# Investigating vaccine uptake dynamics of neutrophils using HIV-1 Envelope glycoprotein trimer

**DOI:** 10.1101/2022.11.10.515906

**Authors:** Philip Y.X. Ngo

## Abstract

Rhesus macaques were previously immunized with two distinct subunit vaccine candidates, to monitor antigen trafficking by immune cells infiltrating the site of injection. The first, a formulation based on the HIV-1 envelope glycoprotein (Env) and the other based on the RSV-fusion (F) protein. Neutrophil and monocyte uptake profiles were vastly different between the vaccine candidates despite similar cell infiltration numbers, hinting that antigen characteristics could orchestrate different innate responses. Notably, the Env trimer is significantly more glycosylated than RSV-F.

Recapitulating these *in vivo* observations under reliable *in vitro* conditions is thus of importance in exploring uptake dynamics and insights into the manipulation of innate responses. The study later demonstrated a complement component, Mannose Binding Lectin (MBL), to be resistant to heat-inactivation and binds Env in a CRDdependent manner. Interestingly, the data suggests a heat-labile component of the serum hinders MBL from binding to Env, which corresponded to a weaker uptake profile. Also, the generation of 3 differentially glycosylated Env variants to study glycan-mediated uptake by neutrophils derived contrary observations. In all, modulation of MBL interactions could potentially target specific innate immune cells, particularly neutrophils, and the later development of adaptive immune responses after immunization.

## Introduction

This project aims to characterize the interactions between a selected HIV-1 envelope trimer vaccine (Env), and a fundamental cell of the human innate immune system through evaluations of phenotypic differences and antigen uptake using flow cytometry. As the topic of innate cell subsets contributing to successful immunization remains mostly unexplored, elucidating the potential roles of neutrophils; the largest constituent of the innate immune system, is of great interest. Thus, this project serves to coalesce queries in Vaccinology surrounding cells of the innate response, such as neutrophil biology and mechanics of antigen uptake.

### Acquired immunity and challenges posed during immunization

Acquired immunity is conventionally achieved through the process of recovery from infectious diseases. Vaccination intends to confer a similar degree of protection towards an agent, but without harm to the host usually through the introduction of protein antigens or attenuated versions of the pathogen (Lombard, Pastoret, & Moulin, 2007; Plotkin, 2014). Consequently, the generation of an adaptive immunity is achieved, capable of long-term memory and recall responses upon future exposure to the immunogen or pathogen (Plotkin, 2014). Attempts in the development of a vaccine against HIV-1, to mitigate Acquired Immunodeficiency Syndrome (AIDS)-related death have been challenging and unrelenting since the 1980s (UNAIDS, 2004). The challenge is partially due to the high degree of variability and mutagenicity of HIV-1 in face of an immune response (Barouch, 2008). Moreover, the functionally crucial epitopes on the target Env spike protein, necessary for the viruses’ ability to bind and enter CD4^+^ cells are obscured by the heavily glycosylated protein surfaces (Cao et al., 2017; Reitter et al., 1998). These challenges epitomize the need for vaccine research to understand the fundamental processes underlying the development of long-lasting and effective immunity (Cao et al., 2017). Modulating aspects of the immune response against the aforementioned HIV-1 characteristics is thus of particular interest to improve immunizing efficacy (Ingale et al., 2016; Zhou et al., 2017).

### Initial immune responses upon immunization, the innate immunity

Most vaccine formulations are commonly administered through i.m injections, where few immune cells predominantly reside in. An adequate delivery of the vaccine antigen to nearby draining lymph nodes (dLNs) would thus rely on rapid immune cell infiltration likely through the events of inflammation, at the site of administration (Seya, 2010; Sun et al., 2009). Liang et al. demonstrated the importance of the early immune responses during immunization, through the generation of different immunization outcomes from the varying magnitudes of initial responses in rhesus macaques (RMs). It remains however, a topic of speculation on the mechanistic principles of these augmented vaccine responses tailored by different immune infiltration kinetics. Differential antigen uptake profile by infiltrating cell subsets is noteworthy, particularly with the dominant cell subset infiltrating the site of immunization, being neutrophils (Liang et al., 2017; Vono et al., 2017; Ols et al., unpublished).

### The initial responder cell: Neutrophils

Neutrophils are the most abundant type of granulocytes, representing a significant fraction of the innate immune system approximating to 60-70% of all leukocytes in circulation of a healthy individual (Amulic et al., 2012; Witko-Sarsat et al., 2000). Expressing a diverse range of pattern recognition receptors such as Toll-like receptors (TLRs), fragment crystallisable (Fc) receptors, and complement receptors, neutrophils swiftly react to events of tissue damage or infection and are rapidly recruited to the site of inflammation through chemotactic responses (Takashima & Yao, 2015). Likewise during immunization modelled in mice, sheep, and RMs, neutrophils are often found in majority at the site of vaccine administration (Calabro et al., 2011; De Veer et al., 2013). To address the perceived threat, neutrophils possess an array of responses such as degranulation, free radical synthesis, the release of neutrophil extracellular traps (NETs), and most notably phagocytosis (Amulic et al., 2012).

### Contemporary understanding of neutrophils during immunization

Neutrophils are often described with myopic lifespans, likely an adaptation to mitigate phagocytised pathogens from parasitizing. There is, however, increasing evidence pointing towards the neutrophil’s unanticipated viability, incorporating antigen-presenting capabilities under certain conditions both during and post-infection (Takashima & Yao, 2015; Vono et al., 2017). In recent literature, Vono et.al demonstrated the antigen-presenting functions of freshly isolated human neutrophils albeit at a lower extent when compared to dendritic cells and monocytes.Neutrophilsl nevertheless have a crucial role in trafficking antigens to lymphatic compartments (Duffy et al., 2012; Maletto et al., 2006)

### Preliminary work by the Loré group: Neutrophil interaction with antigens likely modulated by vaccine characteristics rather than adjuvants

A vaccine-tracking experiment recently conducted in RMs drew interesting observations from the characterization of infiltrating innate immune cell subsets. The majority cell subset that was antigen-positive; an indication of uptake, shifted significantly between neutrophils and monocytes depending on the antigen used (Fig. C). Interestingly, antigen uptake of the monomeric HIV-1 Env gp120 is dominated by neutrophils (Liang et al., 2017), while RSV-F was found to be dominated by monocytes (Ols et al., unpublished). The total number and proportions of the infiltrating immune cells were not any different since similar adjuvants were used. Interestingly, there are more N-linked glycans particularly of the mannose type, found on the surface of Env compared to RSV-F (Gorman et al., 2014).

### Innate immune cells target glycosylated particles to germinal centres

Tokatlian et al. elegantly demonstrated innate immune recognition of glycans and shuttling of nanoparticles to dLNs in a glycan-mediated manner. Follicular Dendritic Cell (FDC) localization, indicative of positive vaccine responses, was significantly abrogated in either MBL-deficient mice or through the deglycosylation of the immunogen, ultimately influencing antigen-specific responses (Tokatlian et al., 2019). The study thus hints at the ability to direct immune responses towards arbitrary particulates, such as vaccine antigens, through manipulation of glycan profiles and therefore its subsequent interactions with the cells of the innate immune system. With each subunit of the Env trimer arrayed with ∼28 N-linked glycans (Cao et al., 2017), the paradigm of how we previously perceived the glycan shield as only impeding effective immune responses (Zhou et al., 2017), also includes it as a target that could amplify immunization outcomes through complement-dependent interactions, specifically MBL (Jack et al., 2001; Lam et al., 2014; Loke et al., 2016).

### Complement system cascade and MBL

The complement system constitutes an archaic but crucial component of innate immunity that ultimately cascades to the formation of the membrane-attack complex (MAC), an unspecific yet effective first-line of defence against most pathogens (Freeley et al., 2016; Merle et al., 2015). Three well-defined activation cascades; the classical, alternative, and the lectin pathway, share many common serum components but differ in the trigger mechanisms. Of particular interest, the members of the lectin pathway compose of a variety of ficolins, collectins, and lectins (Jack et al., 2001; Vorup-Jensen et al., 2014). Although described to exist in various oligoforms, the structure of MBL commonly consists of a carbohydrate-recognition domain (CRD), a collagen-like domain, and a cysteine-rich N-terminal domain (Teillet et al., 2014; Thielens et al., 2014). As a C-type Lectin, recognition of carbohydrate ligands by CRD of MBL is calcium-dependent. The subsequent cascade with the natively conjugated MBL-associated serine proteases (MASPs) are also calcium-dependent (Thielens et al., 2014). In circulation, native MBL complexes with isoforms of MASPs collectively activates C4 and C2 to trigger the complement cascade.

Kuhlman et al. have demonstrated that MBL alone, opsonized the bacteria *Salmonella montevideo*, and facilitated MBL-dependent uptake by neutrophils. These results reinforce the multi-faceted roles of MBL, functioning as an opsonin, and promoting phagocytosis in the absence of other complement components such as MASPs (Brouwer et al., 2014; Kerrigan & Brown, 2009; Kuhlman, 2004).

## HYPOTHESIS AND OBJECTIVES

The importance of a robust innate immune cell infiltration at the site of immunization has been demonstrated to increase the priming of antigen-specific CD4+ T cells in dLNs (Liang et al., 2017). Adjuvant-driven immune cell infiltration and uptake likely promoted efficient antigen delivery to augment T follicular helper (Tfh) cell differentiation and the formation of germinal centres (GCs). Characterization of the muscle-infiltrating cell subsets of 2 vaccine candidates, HIV-1 Env and RSV-F, exhibited distinct uptake profiles that shifted between neutrophils and monocytes being in the majority respectively despite similar infiltration numbers. This study hypothesizes that antigen characteristics, particularly glycan profiles, mediate preferential uptake by distinct subsets of innate immune cells. It is currently not known whether skewing uptake towards different cell subsets would enhance immunization outcomes. An *in vitro* investigation of the uptake dynamics of common infiltrating cells, particularly neutrophils, could provide preliminary insights into the feasibility of manipulating the initial immunization response.

Thus, the project aims to clarify the following:

1. Establishment of a reliable *in vitro* platform to study uptake dynamics of freshly isolated leukocytes.
2. Delineating possible differences in Env uptake profiles between monocytes and neutrophils *in vitro*.
3. Investigating effects of serum components and interactions on Env uptake by neutrophils.
4. Manipulation of glycan-mediated uptake processes in neutrophils.

## MATERIALS AND METHODS

All steps were performed at room temperature unless specified. All wash steps were performed at 1500RPM for 6 minutes unless specified.

### Donor consent

The study was performed in accordance with the Helsinki declaration and approved by the local ethical review board at Karolinska Institutet, Stockholm, Sweden. Buffy coats and whole blood samples were collected from healthy human individuals after informed consent.

### Isolation of human blood neutrophils by Polymorphprep

Human peripheral blood neutrophils were isolated from whole blood collected in either Heparin sodium vacutainer (BD, Sweden) or Hirudin tubes (Sarstedt, Sweden). The blood was processed in sterile conditions in a class II biosafety cabinet as soon as possible upon acquisition, no longer than 1 hour. Polymorphonuclear leukocytes (PMNs) were isolated accordingly with Polymorphprep (Axis-Shield) gradient centrifugation. 3mL of whole blood was layered gently onto 5mL of Polymorphprep solution in a 15mL falcon tube and centrifuged at 2500rpm for 25 minutes without brake and minimal acceleration. The second interface layer from the top is then isolated with a Pasteur pipette into another tube where it was washed with Hank’s Buffer Salt Solution (HBSS) without calcium by resuspension and centrifuged at 1500rpm for 5 minutes, removing the supernatant.

The cell-pellet is then subjected to 1-2 instances of red blood cell lysis when necessary. RBC lysis is performed by resuspension of the cell pellet with 1 part 0.2% NaCl-solution at room temperature for exactly 40 seconds, before supplementation with an equal part of 1.6% NaCl-solution.

The processed cell-pellet is then resuspended in R10 media containing RPMI1640, 1% L-Glutamine, 10% FCS, and 1% Penicillin/Streptomycin (R10 reagents from Hyclone, USA). Isolated PMNs were then quantified with an automated cell counter (Invitrogen) following trypan-blue staining.

The purity of neutrophils isolated was ≥ 70% as indicated in the manufacturer’s protocol and confirmed with CD11b and CD15 staining using flow cytometry (BD Fortessa).

### Isolation of Monocytes from buffy coat (BC) or whole blood

Monocytes isolated from whole blood were processed in parallel with PMN isolation from Polymorphprep (Axis-Shield) gradient centrifugation by removing the first interface layer. The interface was then washed twice with HBSS and resuspended in R10 media. The purity of the monocytes isolated from Polymorphprep is around 10% as confirmed by CD14 and HLA-DR staining under flow cytometry.

Monocytes isolated from BCs were obtained first by enrichment of the BC with 15µL of RosetteSep human monocyte enrichment cocktail (Stemcell Technologies, Sweden) per mL of BC. Followed by layering one part enriched-BC onto two parts Ficoll-Paque (GE Healthcare, Sweden) and then by gradient centrifugation at 2200rpm without brake and minimal acceleration for 20 minutes. The interface was then collected and washed twice with PBS. RBC lysis was performed if necessary. The processed cell pellet was then resuspended in R10 and quantified using the cell counter after staining by trypan-blue.

### Antigen uptake setups

The HIV-1 native flexibly linked (NFL) Env trimer from the South African clade C1086 labelled by Alexa Fluor 680, and the liposomal-conjugated variant (Env-Liposome) was provided by our collaborators from Richard T. Wyatt’s group at the Scripps Research Institute (CA, USA). Env formulations were used at 5μg/ml in each condition unless indicated. RM plasma was obtained from both immunized (24-hours post-immunization with Env-Liposome) and naïve subjects. Human ab serum (Sigma, Sweden) was purchased and tested negative towards HIV-1 specificity as indicated in the batch reports. Heat-incubation of serum/plasma was performed by incubation at 56°C for 30 minutes to inactivate the complement.

The Env trimer used was pre-incubated with 10μL of serum/plasma, Matrix-M™ (Novavax, Sweden) at 0.75μg/ml unless specified, supplemented to a total volume of 100μL with R10 media for 1 hour. Env-Liposome setups were not supplemented with Matrix-M™. Following pre-incubation, 1 million isolated neutrophils or monocytes in 100μL were introduced resulting in a total reaction volume of 200μL. D-Mannose (Sigma, Sweden) was supplemented at 100mg/ml when necessary. Setups were incubated at 37°C with 5% CO_2_ for 24 hours unless specified.

### MBL binding and detection by ELISA

All conditions were run in duplicates and a mean OD value was recorded. ELISAs were performed using Maxisorp plates (Thermo Scientific, Sweden) coated with 200ng of Env variant diluted in PBS overnight. Plates were washed six times with PBS supplemented with 0.05%-Tween-20 between each step. The plates were blocked for two hours using 0.1M CaCl and 1% BSA in PBS (block buffer). Recombinant human MBL (R&D systems) was reconstituted in PBS and serially diluted from 10μg/ml by two-fold dilution. Serum/plasma was diluted 1:10, 1:100, or 1:1000 in block buffer. EDTA was supplemented at 20mM per condition and dMannose at 100mg/mL when specified. The primary antibody, Mouse anti-human MBL monoclonal IgG2a (R&D systems, USA) was applied and incubated for two hours. The secondary antibody, anti-mouse IgG HRP-conjugated antibody (R&D systems, USA) was applied and incubated for one hour. TMB substrate (Biolegend, USA) was added, and the reaction was stopped by the addition of 1M sulphuric acid. MBL binding was quantified using an ELISA Microplate reader (PerkinElmer, USA).

### Deglycosylation of the Env trimer

Env was deglycosylated two ways; demannosylation by EndoH (Promega, Sweden), or complete deglycosylation by PNGase F (New England Biolabs), under non-denaturing conditions according to manufacturer’s guidelines. Deglycosylation of Env was confirmed by SDS-PAGE (Fig. 5) and visualized using Comassie brilliant blue stain. Purification of the deglycosylated variants was performed using size-exclusion ZebaSpin desalting columns (ThermoFisher, Sweden), and Slide-A-Lyzer minidialysis tubes (Thermo-Scientific, Sweden) incubated for 48 hours at 4°C to remove the cleaved glycans.

### Phenotypic characterization using flow cytometry

Configuration of the flow cytometer LSR Fortessa (BD) is indicated in supplementary figure 1. Analysis of Env uptake by neutrophils or monocytes was done using the detection of AF680. Analysis of the Liposome signal on the Env-Liposome vaccine was done using the detection of Top-Fluor signal in the FITC channel. The TopFluor signal is subsequently used as a control for the correct vaccine formulation used when characterizing uptake profiles. All setups were stained with 5μL of human FcR blocking reagent (Miltenyi Biotec, USA) to mitigate unspecific binding and LIVE/DEAD blue stain (Invitrogen, Sweden). Staining panels for monocytes and neutrophils are indicated in the supplementary tables 1 and 2 respectively. Events acquired for each condition exceed 500,000 events unless specified. Gating strategies using FlowJo-10 (FlowJo Inc., USA) for monocytes and neutrophils are indicated in the supplementary figures 1 & 2 respectively.

**Suppl. Fig. 1.**
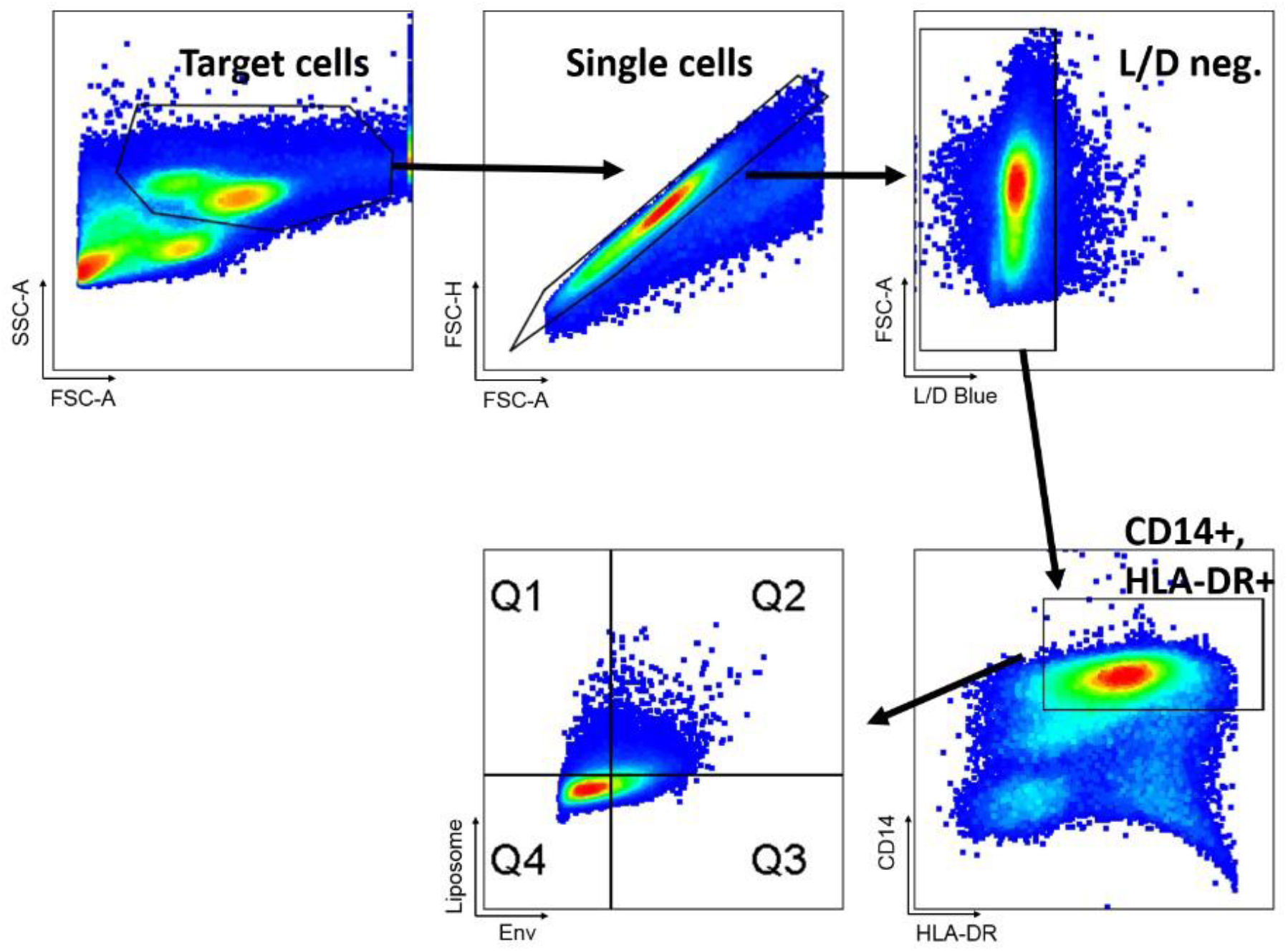
Monocyte gating strategy in quantifying Env+ cells. Q1/2 represents Liposome+ cells, Q2/3 represents Env+ cells. Gating strategy is justified from negative population controls and population morphologies with compensation using FlowJo v10 software.

**Suppl. Table 1.**
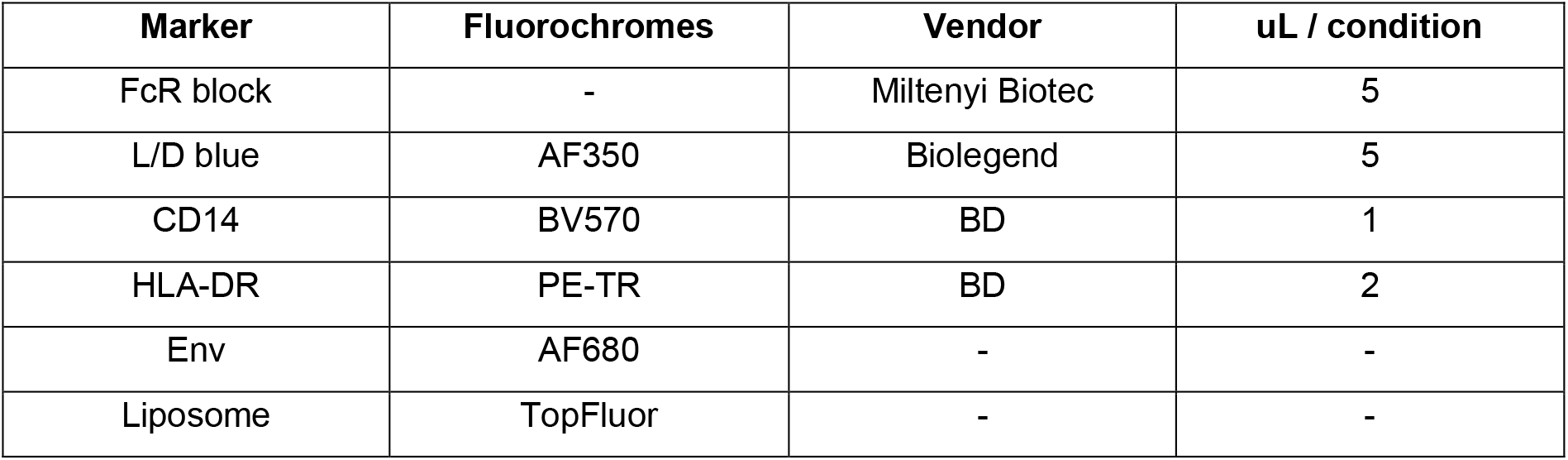
Antibody staining panel and fluorochrome detection for monocyte uptake assays

| Marker | Fluorochromes | Vendor | uL / condition |
| --- | --- | --- | --- |
| FcR block | - | Miltenyi Biotec | 5 |
| L/D blue | AF350 | Biolegend | 5 |
| CD14 | BV570 | BD | 1 |
| HLA-DR | PE-TR | BD | 2 |
| Env | AF680 | - | - |
| Liposome | TopFluor | - | - |

**Suppl. Fig 2.**
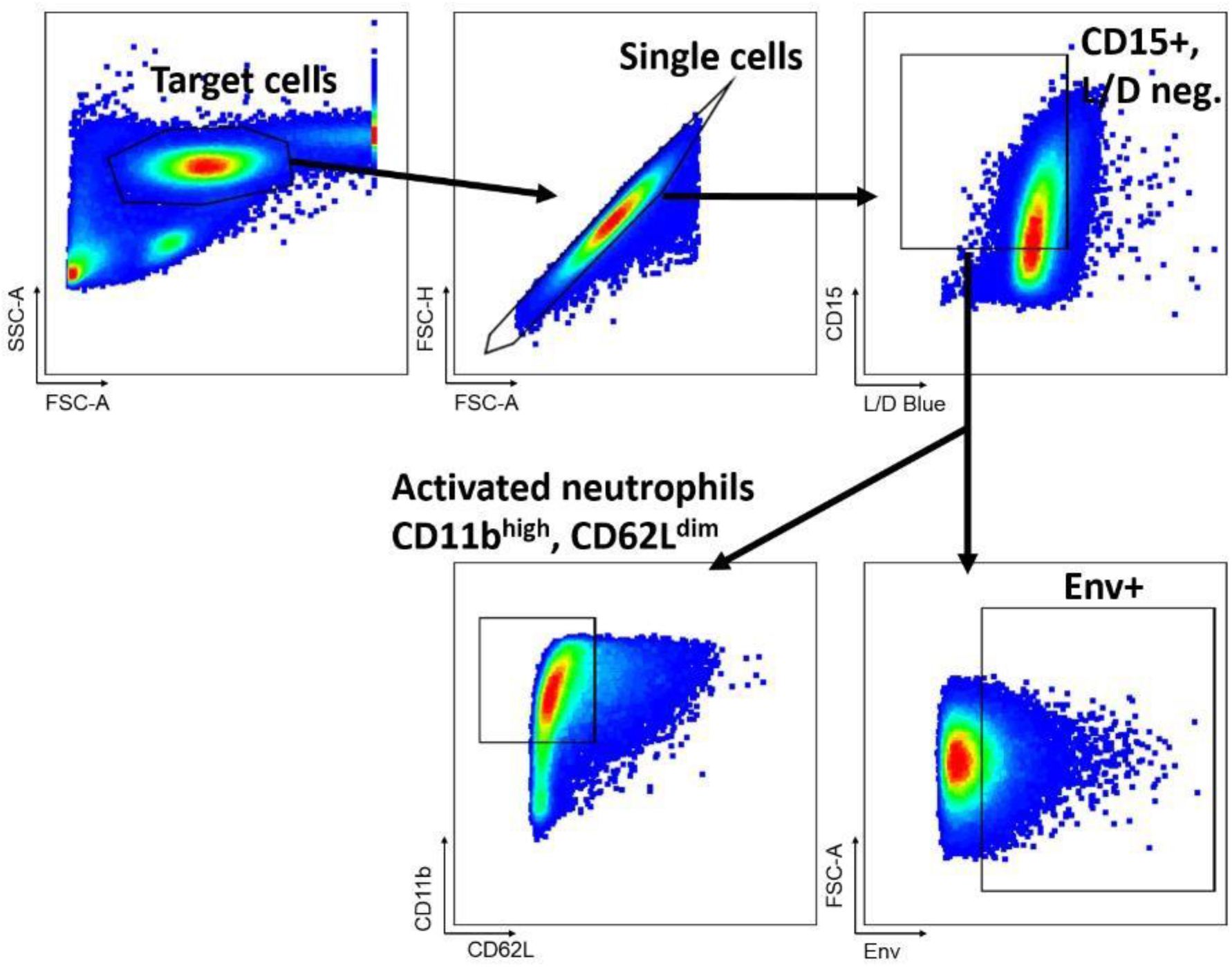
Neutrophil (PMN) gating strategy after processing with PolyMorphPrep. Observations on the activation of neutrophils are optional in the analysis. Gating strategy is justified from negative population controls and population morphologies with compensation using FlowJo v10 software.

**Suppl. Table 2.**
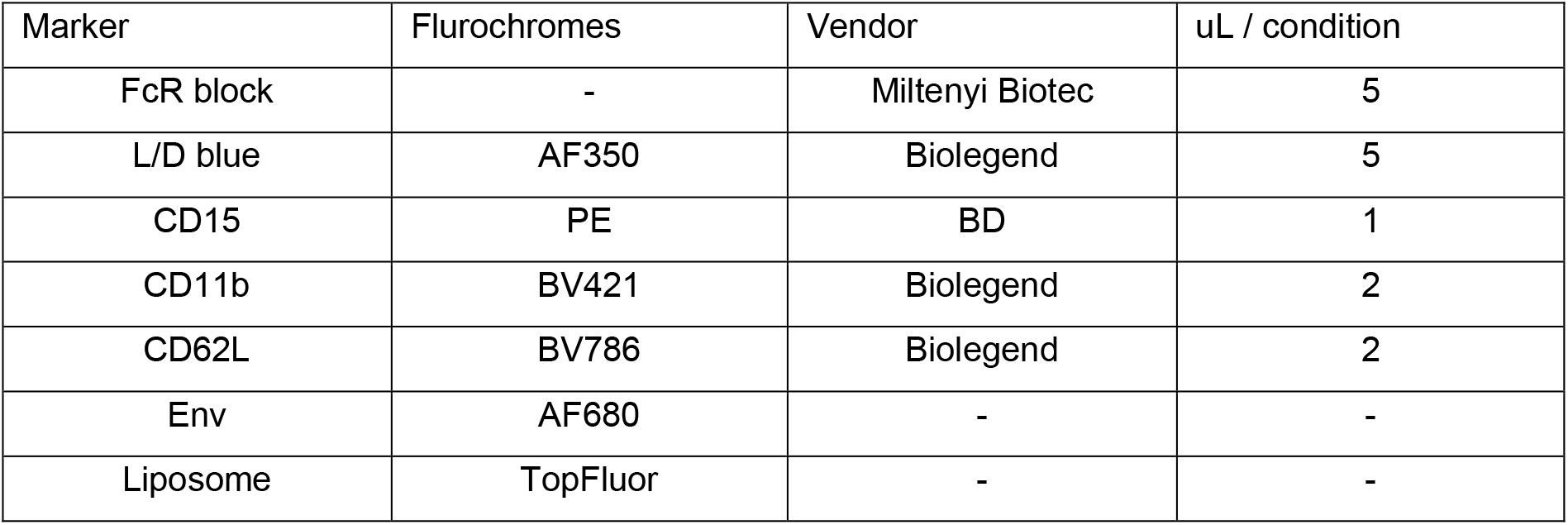
Antibody staining panel and fluorochrome detection for neutrophil uptake assays

## RESULTS

### Validation of the previously established protocol to study antigen uptake *in vitro*

Monocytes were previously demonstrated by the group to possess active uptake response in-vitro, regardless of the immunogenicity of the test molecule. This setup thus serves as a positive indication that the protocol could be extrapolated to other cell types as well. Consistent with previous attempts by other members of the group, monocytes demonstrated observable uptake responses when provided with either soluble Env trimer, or the Env-Liposome formulation (Fig. 1.1, 1.2). Uptake responses were detectable even in the absence of blood plasma (not shown). Uptake responses fell modestly when a heat-inactivated variant of blood plasma was used, suggesting the involvement of complement components in driving the uptake of these formulations (Fig 1.1 A, 1.2 A).

Interestingly, the use of blood plasma from RMs previously immunized with EnvLiposome promoted a substantial uptake response. This was however diminished with the heat-inactivated variant of the immunized plasma (Fig. 1.1 B, 1.2 B).

**Fig. 1.**
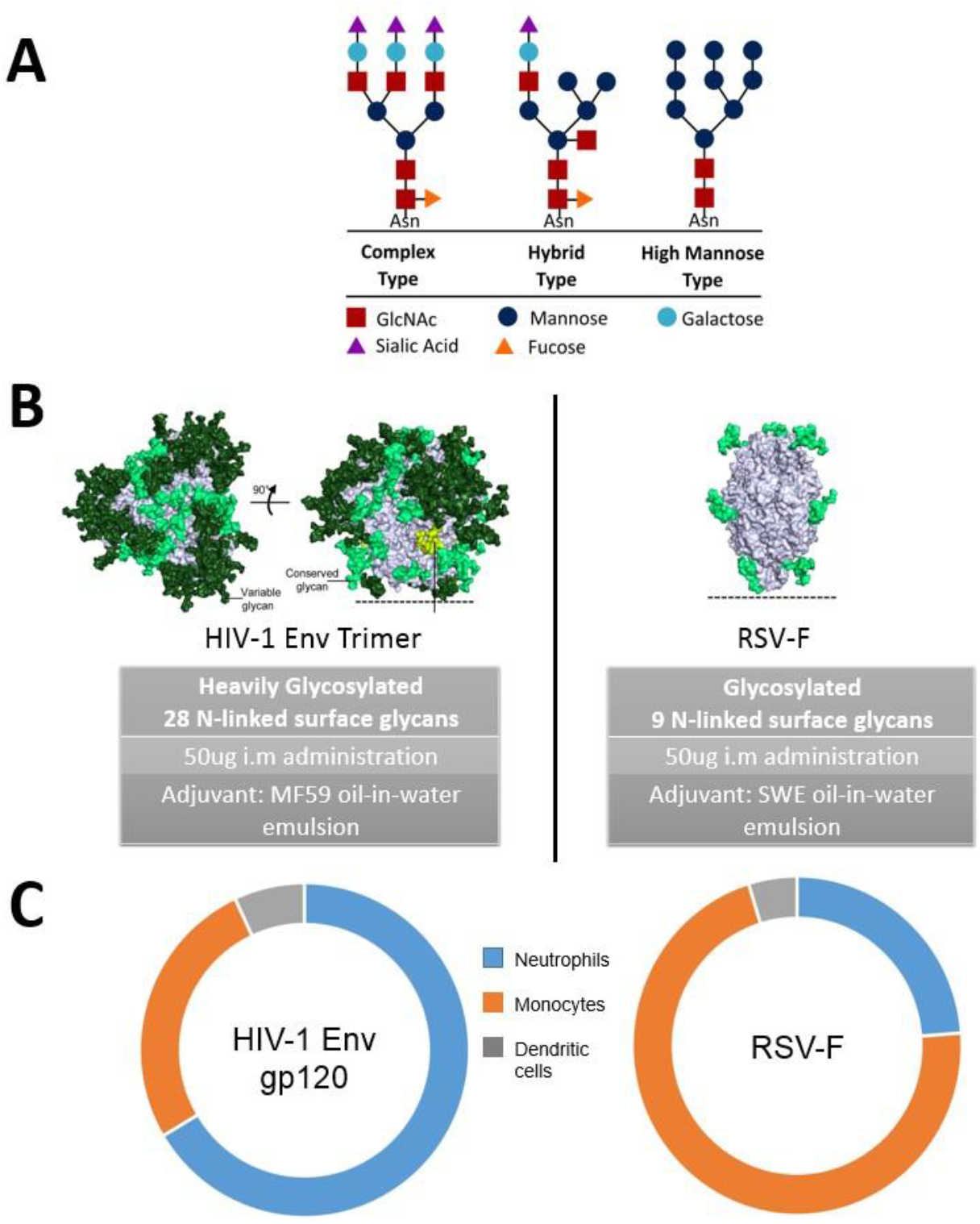
Preliminary work on antigen-trafficking and initial immune cell infiltration. (**A**) The types of glycans found on the vaccine candidates used to immunize the RMs (Liang et al., 2017, Ols et al. unpublished). (**B**) [Left], a HIV-1 Envelope subunit consists of 28 N-linked surface glycans, of which 14 are high-mannose type, 6 are complex type, and 8 variable hybrid types. Env is considered heavily glycosylated as compared to [Right] the RSV-F formulation, consisting of only 9 high-mannose type glycans. Both formulations were administered i.m at 50ug/site, supplemented by similar oil-in-water emulsion-based adjuvants. Image modified from Gorman et al., 2014. (**C**) Pie-chart depicting the subsets of vaccine+ infiltrating cells found at the site of administration 24-hours post-immunization (Ols et al., unpublished).

**Fig.1.1.**
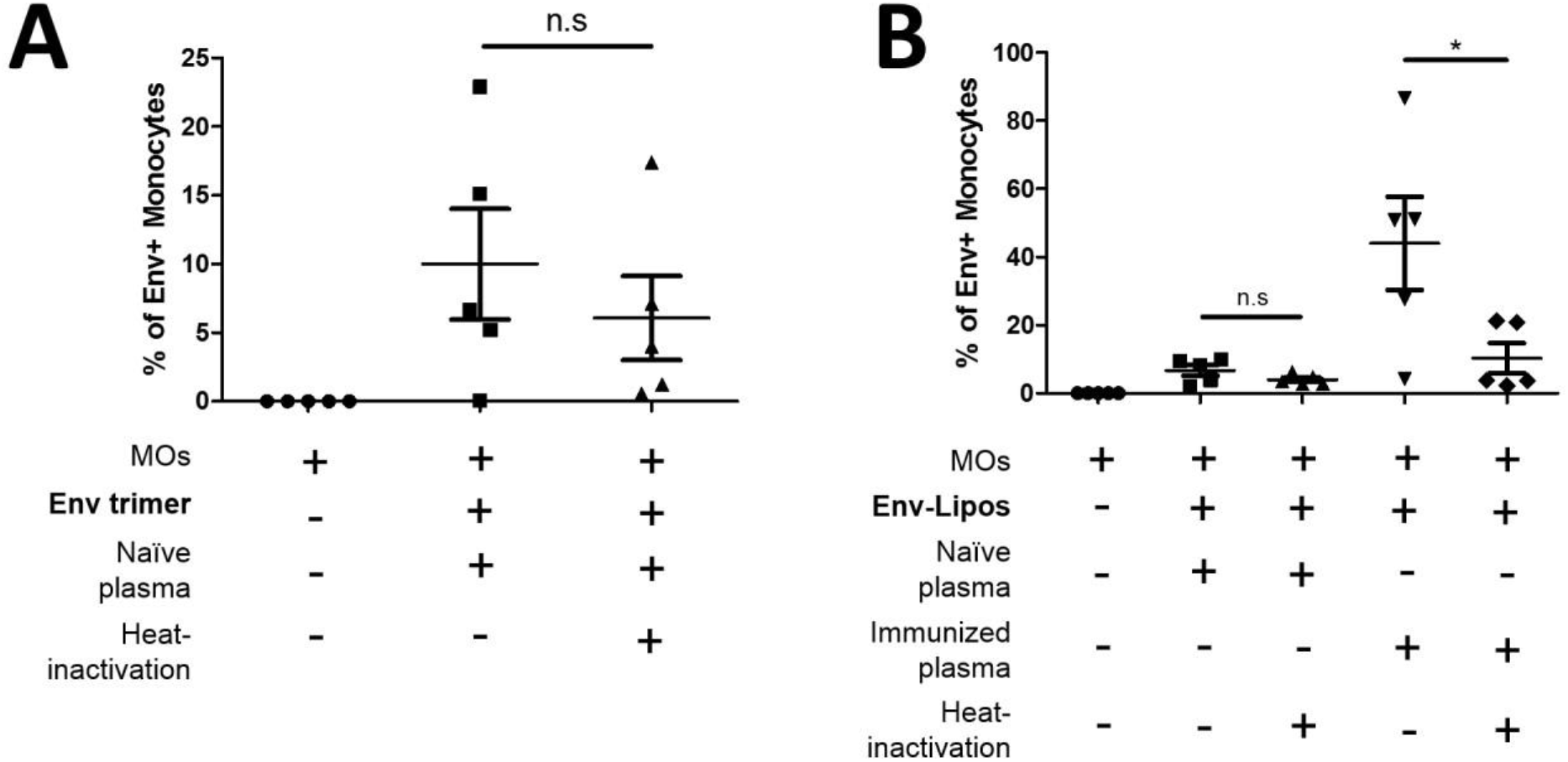
Heat-labile complement components augments antigen uptake in monocytes. Error bars represent SEM. Monocytes were isolated from buffy coats, using the same set of 5 donors for each uptake profile. *P <0.05 (Mann-Whitney) (**A**) Soluble Env trimer uptake profile of monocytes expressed as the number of Env positive cells over total monocytes gated from live, CD14 and HLA-DR double positive cells (n=5). (**B**) Env-Liposome uptake profile of monocytes gated similar to Env trimer uptake, (n=5).

**Fig. 1.2.**
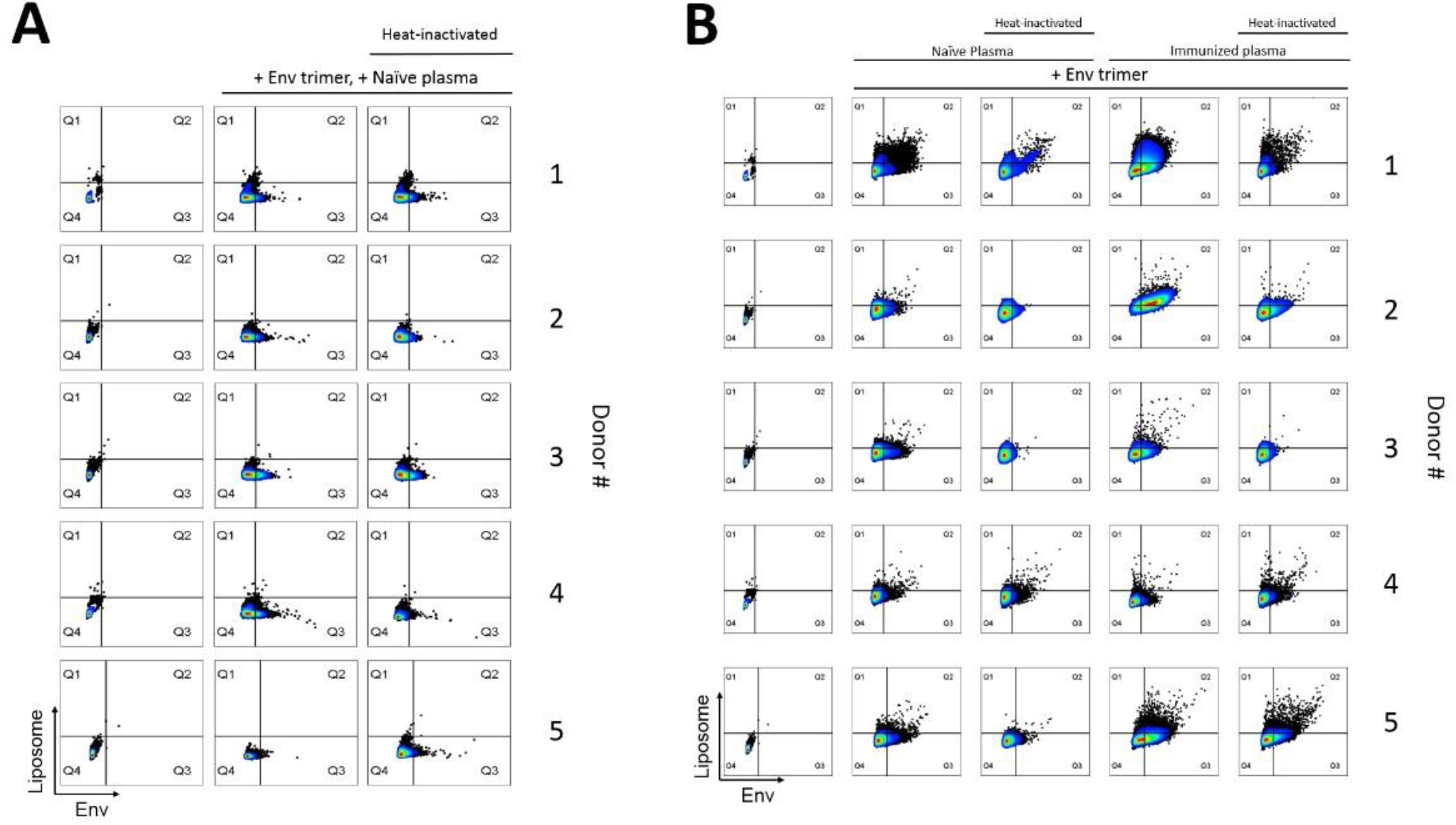
Flow-plots of Env positive monocytes (AF680), against TopFluor signal for Liposome positive cells. (**A**) Soluble Env trimer uptake, (**B**) Env-Liposome uptake. Gated in Q2 and Q4 are quantified as Env+ cells. Gate in Q1 and Q2 are considered as Liposome+ cells.

### *In vitro* uptake of Env by Neutrophils (PMNs) is augmented by heat-inactivation of complement components

Despite Env uptake responses being discernable within an hour of incubation in monocytes, Env uptake by neutrophils was only evident after 24 hours of incubation (Fig. 2.1A). Env-Liposome uptake remains apparent within an hour of incubation however (Fig 2.1B). To try to simulate *in vivo* conditions closely, Matrix-M was included in conditions where Env trimer uptake was investigated. However, the effects of Matrix-M on neutrophil uptake of Env were found to be minimal (Fig 2.2). Interestingly, uptake responses by neutrophils were augmented by the heat-inactivated variants of both types of RM plasma, regardless of the Env formulation used (Fig 2.1 A, B), and contrary to the observations with monocytes. To evaluate if this phenomenon was exclusive to the heat-activated variants of the RM plasma, human ab serum was used instead and a similar albeit modest effect was observed (Fig. 2.1C).

The project now steers towards elucidating potential heat-stable components of the serum that could augment uptake response. Kuhlman et.al previously demonstrated that MBL functions as an opsonin, driving bacteria uptake responses in neutrophils. MBL thus serves as a potential serum component candidate that could influence Env uptake responses in neutrophils.

**Fig. 2.1.**
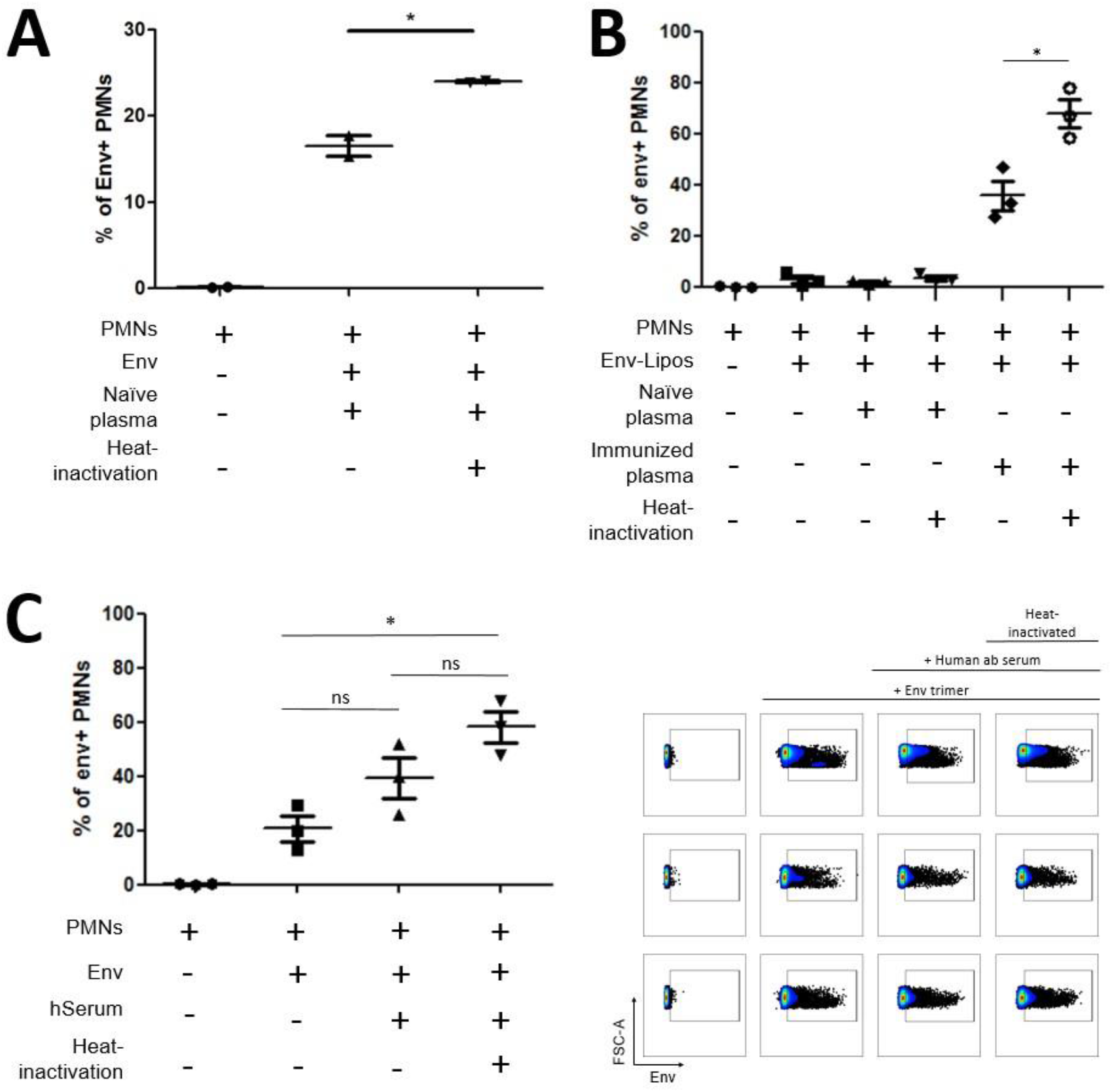
Neutrophils (PMNs) uptake responses with Env trimer or Env-Liposome augmented in heat-inactivated variants of blood plasma from either human or RM derivatives. * Denotes P<0.05 (Mann-Whitney) **(A)** In-vitro uptake profile of PMNs with soluble Env trimer using naïve macaque plasma. (n=2) **(B)** In-vitro uptake profile of PMNs with Env-Liposome formulation using either naïve or immunized macaque plasma. (n=3) **(C)** [Left] Uptake profile of PMNs with soluble Env trimer using human ab serum (Naïve equivalent), [Right] Flow-plots of the uptake profile. Gated are regarded as Env+ cells (n=3).

**Fig 2.2.**
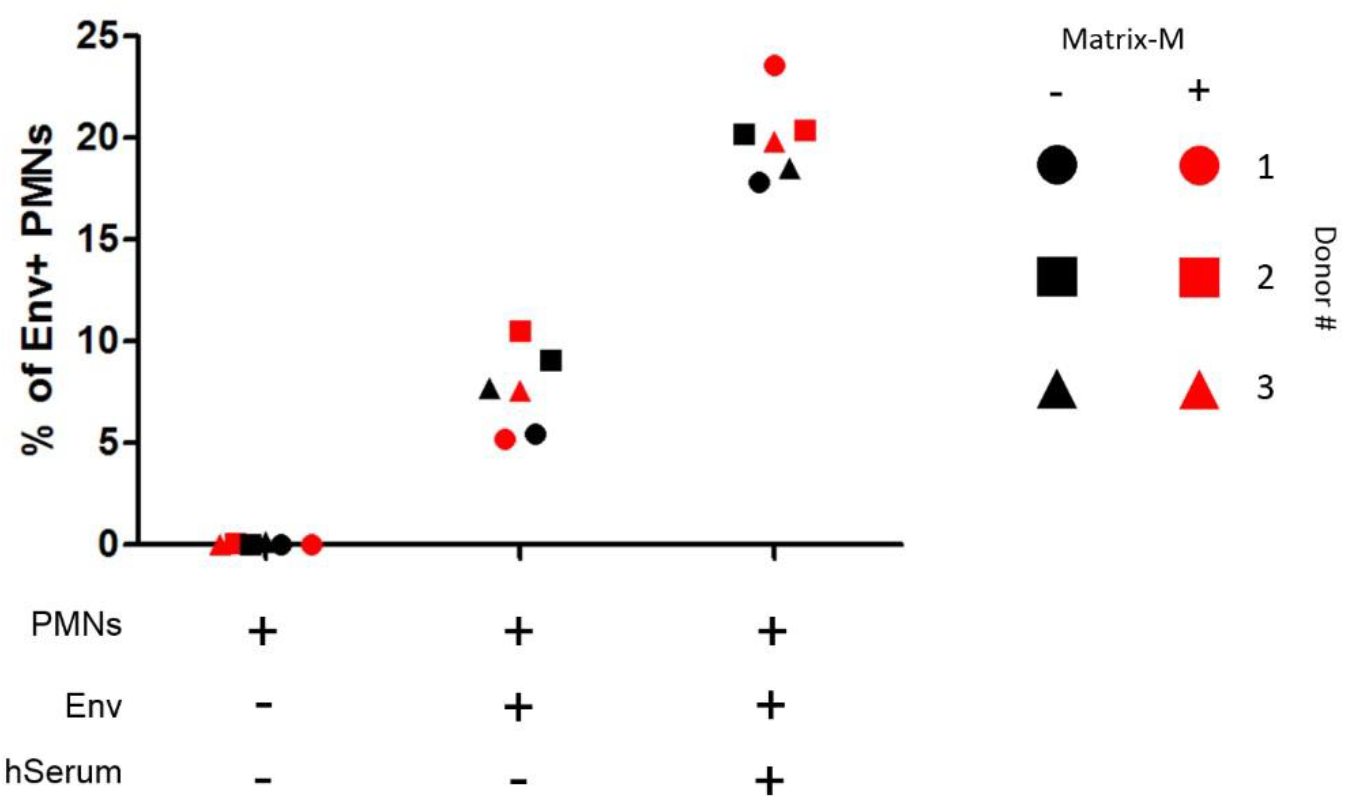
Uptake profiles of neutrophils (PMN) with or without the addition of the adjuvant, Matrix-M.

### MBL detected in serum types, binds to Env trimer in a CRD-dependent manner

Recombinant human MBL (rhMBL) binding to Env coated plates was detected by ELISA (Fig. 3 A, B), and provided a substantial indication that MBL in serum could theoretically opsonize Env. To affirm this hypothesis, rhMBL was substituted with equivalent volumes of human ab serum and the heat-inactivated variant used in uptake assays to assess whether i) MBL was present, and ii) could bind Env. MBL was detected in both serum types (Fig. 3 C) and interestingly, an increase in MBLbinding was quantified in the heat-inactivated human ab serum. This observation aligns with the augmented uptake responses observed with neutrophils in heatinactivated serum types and prompt a plausible correlation between MBL-bound Env, and its uptake.

**Fig. 3.**
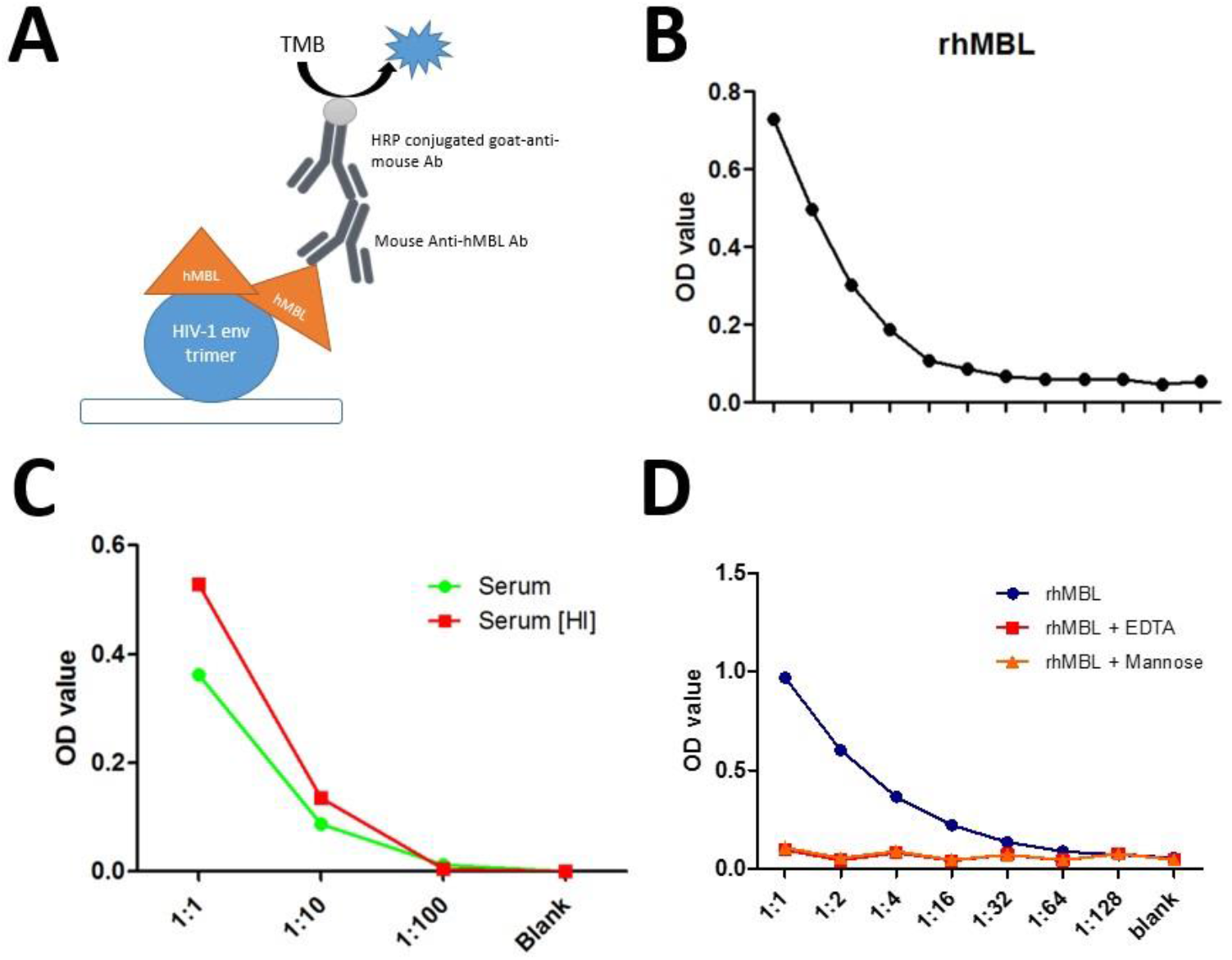
Mannose-Binding-Lectin (MBL) in serum, binds to Env in a CRD-dependent manner. (**A**) Schematic representation of the method, ELISA, used to detect whether human MBL (hMBL) binds to Env. (**B**) Recombinant Mannose-Binding-Lectin (rhMBL) binds to Env. 1000ng of rhMBL was added and diluted 2-folds subsequently as indicated on the X-axis. (**C**) MBL was detected in human ab serum. Equivalent amounts of MBL is assumed between the 2 setups as the serum was split from the same vial. Higher amounts of MBL bound Env is observed with the heat-inactivated serum setup. (**D**) MBL binding to Env is attenuated in the absence of calcium, or in the presence of MBL-CRD ligands, mannose.

An evaluation of whether MBL binding to Env was glycan-mediated was followed up and found that binding of Env was attenuated in the presence of soluble mannose (Fig. 3 D) or when calcium ions were chelated using EDTA. These observations thus demonstrates that MBL binds to the glycans of Env through the carbohydraterecognition-domains (CRD) in a calcium-dependent manner.

### MBL likely opsonizes Env, mediating uptake in neutrophils in the absence of complement components

Upon confirmation by ELISA that MBL detected in serum binds to Env in a CRDdependent manner as represented by rhMBL, recapitulating if the MBL-Env complex could independently influence uptake in neutrophils is thus of interest. Dilutions of rhMBL; 10ng, 100ng, and 1000ng, were added to cultures to investigate a dose-dependent uptake effect (Fig 4. A). Uptake responses were modestly elevated with increasing amounts of rhMBL in culture, with 1000ng of rhMBL significantly augmenting the Env uptake response compared to the control. MBL-mediated uptake was expectedly abrogated in setups containing soluble mannose (Fig. 4. B), demonstrating again that MBL binds specifically to the glycans of Env, and the MBL-Env complex augments the uptake response.

**Fig. 4.**
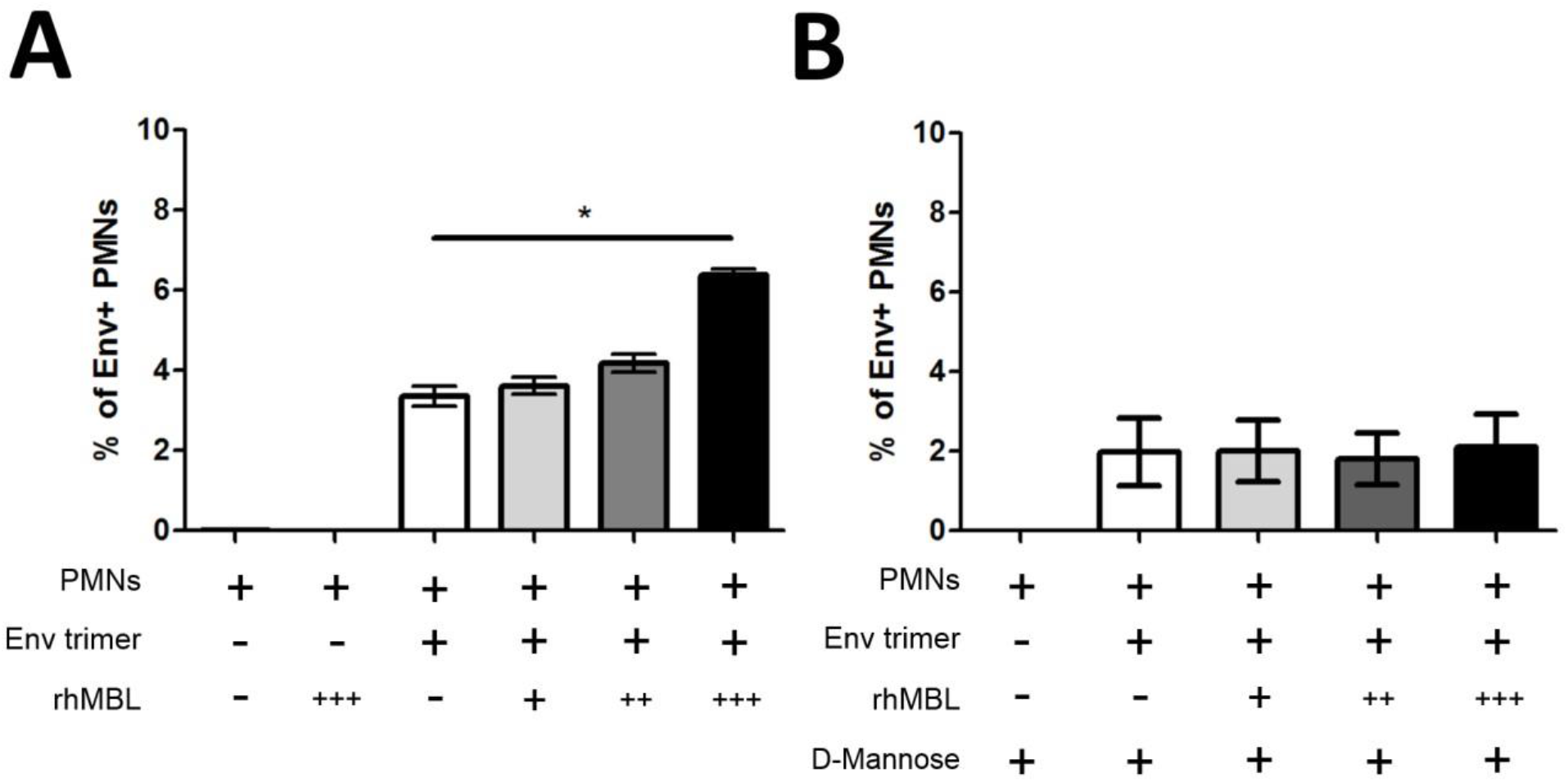
MBL-mediated uptake response in Neutrophils is dependent on CRD binding to Env. *P <0.05, (Kruskal-Wallis). Indications for rhMBL added: +++ represents 1000ng, ++ at 100ng, and + at 10ng. (**A**) MBL directly influences Env uptake by neutrophils (n=3). (**B**) MBL-mediated uptake is absent in the presence of soluble mannose, a known ligand to the CRD of MBL (n=3).

### Generation and in-vitro uptake of demannosylated Env (dgEnv) and completedeglycosylated Env (cdgEnv) by neutrophils (PMNs)

Following the abrogated Env uptake in neutrophils by the introduction of soluble MBL ligands, mannose, it would thus be of interest to verify if this phenomenon could be recapitulated by removing the glycan targets of MBL on Env. The generation of the deglycosylated Env variants was verified by MW shifts on SDS-PAGE (Fig. 5) and purified as indicated in the methods.

**Fig. 5.**
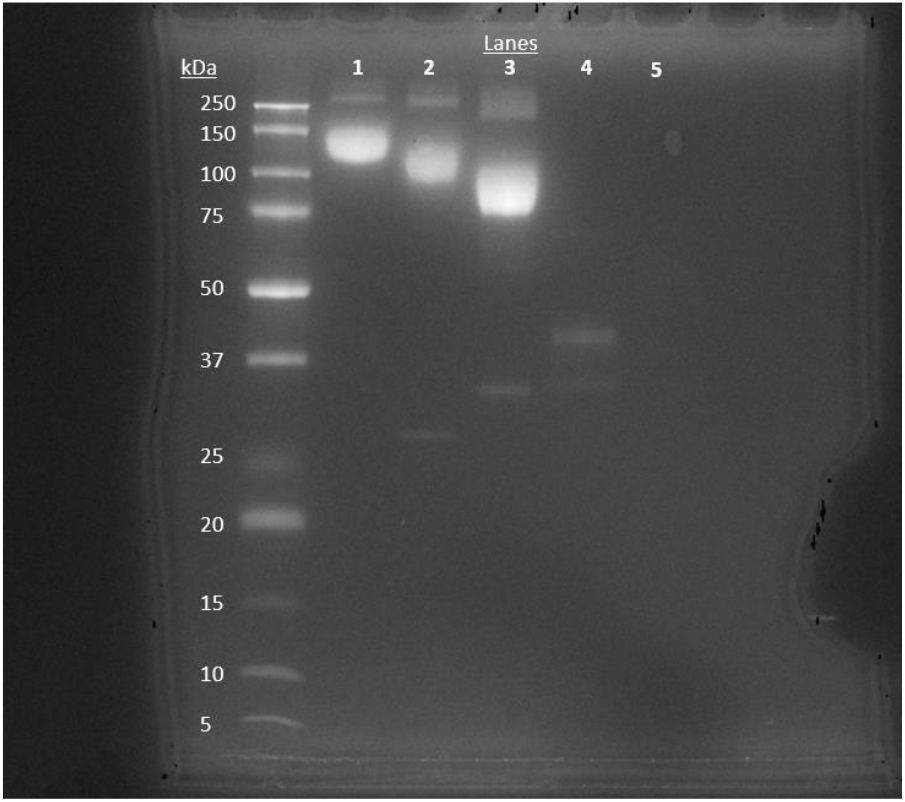
SDS-PAGE check of differentially glycosylated Env. Lane 1 from the left, after the MW ladder, indicates Env that was not subjected to digestion at ∼150kDa. **Lane 2** depicts the demannosylated Env variant by EndoH treatment, further down the lane indicating the presence of the glycosidase at ∼30kDa. **Lane 3** depicts the completely deglycosylated Env variant by PNGase F treatment, further down the lane indicating the presence of the glycosidase at ∼35kDa. Lane 4 and 5 are the positive and negative controls respectively, as supplied with the PNGase F product.

The uptake profile of each Env variant was next assessed with isolated neutrophils and the results were contrary to our predictions. A moderately elevated uptake response was observed for dgEnv, whereas clear inflation of Env+ neutrophils was evident in cdgEnv conditions (Fig. 6 A, B). These observations were consistent when tracked individually of the 4 donors (Fig. 6 A), and were also apparent in a previous pilot of the setup. Setups that were supplemented with human ab serum yielded inconsistent findings between donors but exhibited a moderate increase in overall mean uptake response (not shown) for all Env variants.

**Fig. 6.**
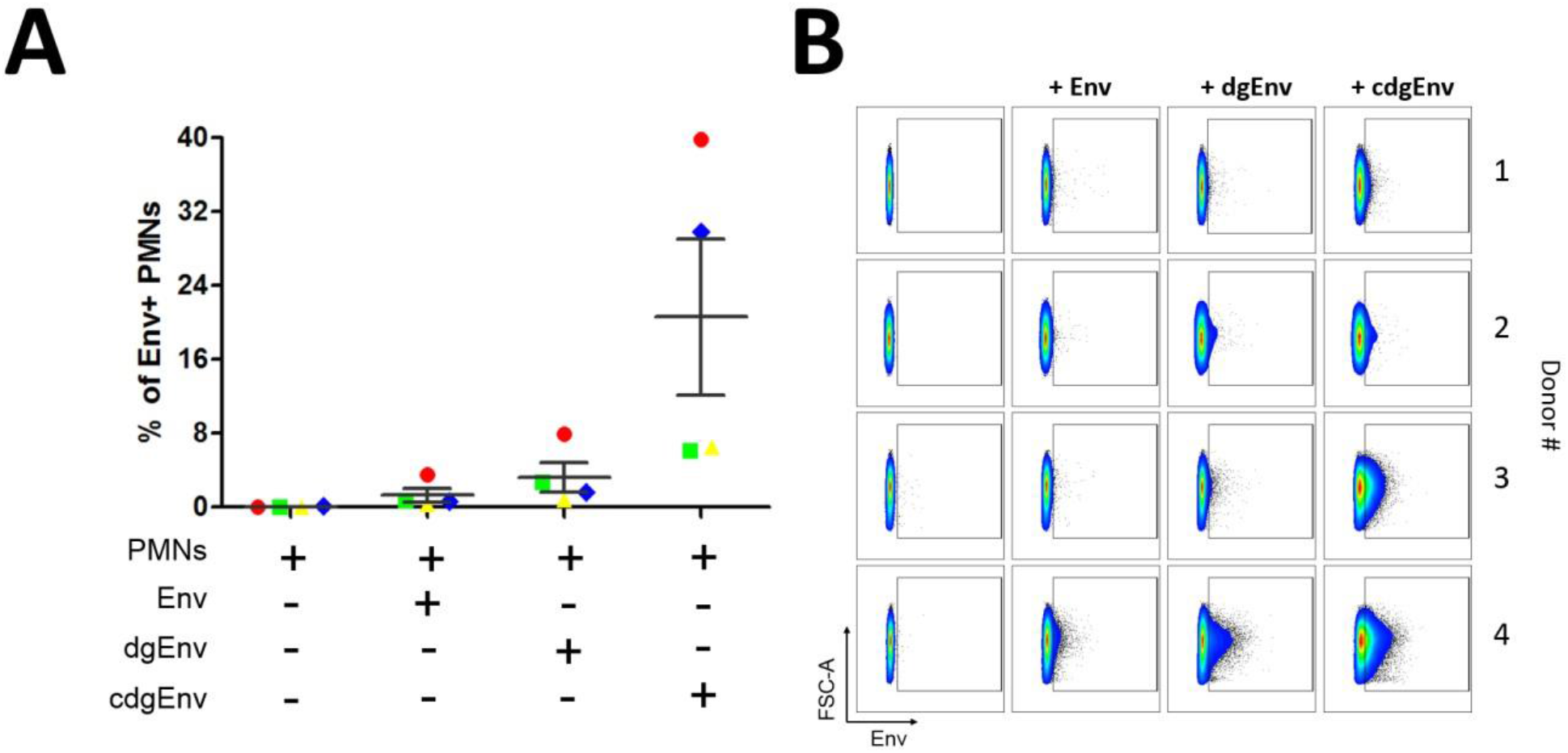
Elevated uptake response in neutrophils with increasing degrees of deglycosylation on Env. (**A**) In-vitro uptake response of 3 differentially glycosylated Env variants with neutrophils. Donor # are indicated with different colors, yellow as #1, green as #2, blue as #3, and red as #4. (**B**) Flowplots of the uptake responses, gated are Env+ cells.

## DISCUSSION

Early innate responses dictating immunization outcomes have long been a topic of speculation. Recently, Tokatlian et al. demonstrated innate immune recognition of glycosylated targets on an HIV nanoparticle, which correlated with enhanced antigen trafficking to GCs in mice. The immunization responses elicited by differentially glycosylated HIV nanoparticles were shown to differ in terms of trafficking and concentration to the GCs. It was noted that trafficking of these nanoparticles was dependent on complement, MBL, and the glycan profile of the immunogen (Tokatlian et al., 2019). However, it is crucial to note that these models in mice possess immune cell populations and innate responses that may differ considerably from humans (Mestas & Hughes, 2004).

RMs however, provide substantial translational value when representing human immunization outcomes albeit a different set of problems not limited to ethical considerations and inter-subject variability; of which is circumvented in studies with inbred mice. Inter-subject variability, however, may provide a more accurate representation of the immunological phenomenon as it accounts for the variations that exist among human individuals (Liston et al., 2016). The Loré group has previously demonstrated that superior vaccine responses are driven by adjuvant-mediated enhancement of innate responder cells and their uptake of antigens (Liang et al., 2017). Later, the group also demonstrated that the major subset of the initial responder cells, neutrophils, can function with antigen-presenting capabilities in regulating specific T-cell responses (Vono et al., 2017). Interestingly, it was also observed during the group’s vaccine-tracking experiments with a novel respiratory syncytial virus (RSV) vaccine candidate that the uptake response was instead comprised of mainly monocytes instead of neutrophils (Ols et al., unpublished). The HIV-1 Env trimer is also found significantly more glycosylated than the RSV vaccine. Under this context, it is thus of interest to delineate the circumstances that influence the uptake of antigens by neutrophils, with a focus on the glycan profile of the vaccine antigen and the serum components present *in vivo*.

### *In vitro* uptake of Env by Monocytes is likely mediated by complement components

Albeit not the focus of this thesis, the Env uptake profiles of monocytes served as an important control when analysing the uptake profiles generated by neutrophils. Interestingly, the improved uptake response elicited from the use of blood plasma from a previously immunized RM suggests that immune complexes (ICs) are likely involved in augmenting antigen uptake responses in monocytes.

ICs are known to trigger an array of responses including opsonophagocytosis, and complement deposition (Goldsby Richard, 2002). The results however suggest that ICs in this case, are likely secondary in driving the elevated uptake response. With diminished uptake responses consistently observed with the heat-inactivated immunized plasma (Fig 1.2 B), it is likely that the ICs formed with the antigen did not promote a direct uptake response from the monocytes but enhanced complement deposition and mediated augmented responses. These results nevertheless reinforced the protocols previously established by the group and demonstrated that uptake of the Env trimer is possible *in vitro*.

### *In vitro* uptake of Env by Neutrophils (PMNs) augmented by heat-inactivation of complement components

The project next examined the antigen uptake dynamics of neutrophils in-vitro and demonstrated improved antigen uptake in the presence of serum components. The uptake response was further augmented when the complement components were heat-inactivated.

These results were contrary to our previous observations of diminished uptake with heat-inactivated serum types, suggesting a heat-stable component of the serum motivating the augmented uptake response of neutrophils but not monocytes. Activation markers of neutrophils (Fortunati et al., 2009; Mann & Chung, 2006); CD62L, and CD11b, were tracked and found unchanged between all setups, which suggests that it is unlikely that the heat-inactivated serum type was more immunogenic, driving the augmented responses. The effects of Matrix-M (Sun et al., 2009) were also evaluated but not influencing uptake profiles significantly (Fig. 2.2).

Kuhlman et.al previously demonstrated MBL functioning as an opsonin in the absence of complement components, augmenting the uptake of the opsonized bacteria in neutrophils. We thus narrowed down our investigation of heat-stable compounds and included the criterion that the serum component should be able to function outside of the heat-labile complement cascade. MBL fits these criteria and we, therefore, investigated the likelihood of MBL orchestrating Env uptake in neutrophils.

### MBL detected in serum, binds to Env trimer in a CRD-dependent manner

MBL consists of a C-terminal calcium-dependent carbohydrate-recognition-domain (CRD), a collagenous stalk, and a cysteine-rich N-terminal domain. Ligands of the CRD in MBL are not limited to mannose sugars, but fructooligosaccharides and to some extent, chitosan as well (Kjærup et al., 2014). Analysis by ELISA revealed that MBL was present in serum, bound to the glycans of Env in a CRD-dependent manner and interestingly, the binding response was augmented likewise when complement components in the serum were heat-inactivated. The amount of MBL is likely similar between the setups as the heat-inactivated variant derived directly from splitting the vial containing the native human ab serum used. These results thus suggest the possibility of a heat-labile component in the serum inhibiting MBL from binding to Env through its CRD regions.

A previous study has demonstrated a dose-dependent inhibition of MBL binding to cell receptor Cr1, by the complement component C1q (Ghiran et al., 2000). It is however not known if complement components, outside of other lectins, could directly inhibit MBL interactions with its ligands. Recapitulating if MBL-Env interactions could directly modulate Env uptake in neutrophils would be next of interest.

### MBL likely opsonizes Env, mediating uptake in neutrophils in absence of complement components

The results demonstrated that MBL can independently influence Env uptake in neutrophils in a dose-dependent manner. With reference to the ELISA data generated previously, it is likely that MBL binding to Env may have been more efficient in promoting uptake in neutrophils than with conventional complement components in the serum. The addition of soluble mannose into a replicate setup abolished the augmented uptake previously driven by rhMBL. These observations thus recapitulated that MBL complexes with Env in a CRD-dependent manner and demonstrated MBL augmenting Env uptake in neutrophils in the absence of complement.

It would next be of interest to understand if this MBL-Env complex can be disrupted similarly if we removed the potential ligands of MBL on the Env trimer.

### Generation and *in vitro* uptake of demannosylated Env (dgEnv) and completedeglycosylated Env (cdgEnv) by neutrophils (PMNs)

The project next generated differentially glycosylated Env variants through glycosidase digestion with either EndoH (dgEnv), or with PNGase F (cdgEnv). The variants were purified using dialysis and size-exclusion desalting columns to remove the cleaved saccharides from the Env variants. The glycosidases were previously mentioned to cease function upon depletion of the reaction buffers, which was achieved during the dialysis process but however, not purified out from both Env variants. With the enzymatic potential of the glycosidases presumably abrogated, the project proceeded with testing the uptake profiles of both dgEnv and cdgEnv in neutrophils. Interestingly, even in the absence of serum components, there was already noticeable augmentation of uptake with the deglycosylated Env variants dgEnv and cdgEnv, when compared to intact-glycosylated Env (Fig.6. B).

In hindsight, it would have been appropriate to analyze the structural integrity of these Env variants using dynamic light scattering or TEM before proceeding. Although the Env trimer did not fragment into monomers upon deglycosylation, as visualized on SDS-PAGE, the subunits were likely held together due to the flexible peptide covalent linkages (NFL) as Wyatt and colleagues previously have described. Treatment with glycosidases may have instead disrupted the stability of the trimer, potentially linearizing the molecule and exposing various epitopes previously concealed by the native quaternary conformation. In the preceding case, it would have resulted in an unfair comparison between the deglycosylated variants as there is more than one variable altered. It is thus necessary to keep in mind these considerations when interpreting the results of this section.

## Future directions

### MBL depletion of serum types

Depletion of MBL in serum using mannose-specific bead separation columns may be of interest to delineate other serum components that may have augmented the uptake response in heat-inactivated serum. As the glycan profile of Env extends beyond heavy mannosylation, it is equally reasonable that other lectins associating with other saccharide groups on Env could have augmented the uptake response in neutrophils as well.

### Dissecting inhibitory components in serum on MBL interactions with Env

ELISA is a technique to demonstrate avidity and affinity of MBL with Env. Dilutions of whole serum or serum components such as C1q or C3, can be added with MBL in Env coated plate to assess the changes in MBL-Env interactions. In addition, the avidity of MBL-Env can be verified using sodium thiocyanate dilutions. The results can later be recapitulated with uptake assays *in vitro* to dissect the different serum components contributing to Env uptake in neutrophils.

### Structural verification of dgEnv/cdgEnv by light scattering and TEM and further purification

Dynamic light scattering and transmission electron microscopy (TEM) are common techniques employed in other studies (Lorber et al., 2012; Sharma et al., 2015; Tokatlian et al., 2019) to verify the structural integrity of a purified compound. Purification of the Env variants from the glycosidases and the cleaved saccharides can be performed through size-exclusion using high performance liquid Chromatography (HPLC) to elute the different Env variants. The purified compound can then be assessed by dynamic light scattering or TEM, to verify the structural stability after deglycosylation. Alternatively, saccharide removal and the successful isolation of the deglycosylated Env variants can be verified on SDS-PAGE with a glycoprotein stain.

### In-vivo tracking of purified dgEnv and cdgEnv in RMs with vaccine outcomes

Referencing the study by Tokatlian et al. and harnessing the techniques of the Loré lab, it would be interesting to recapitulate the observations in RMs. Alternatively, it would also be of interest to investigate if inhibition of MBL-Env formation at the injection site with the addition of mannose in mice would produce any effect on infiltration and the generation of an antigen-specific response.

## CONCLUSIONS

The results have reinforced previous findings of MBL functioning in the absence of complement and provided insights on the interactions between MBL, the HIV-1 Env glycoprotein trimer, and neutrophils in-vitro. The study has demonstrated that MBL binds to Env in a CRD-dependent manner and that MBL-Env augments the uptake profile of neutrophils. The observations further hint at MBL-Env interactions driving superior uptake than conventional serum components, while also suggesting the plausible attenuation of this augmented effect *in vivo* by supplementing MBL ligands. Manipulation of Env glycan profile proved to be challenging but nonetheless a promising avenue to recapitulate the strong influences of the glycan-mediated process on the uptake in neutrophils thus far.

Successful *in vitro* investigations on uptake dynamics of initial responder cells provide firsthand insights into the feasibility of such manipulation *in vivo*, tailoring uptake of antigens from specific cell subsets to generate an overall stronger immunization outcome. In all, this study has demonstrated the potential of MBL in serum influencing the uptake responses in neutrophils of heavily glycosylated antigens, such as Env.

## ABBRIEVATION

AIDS: Acquired Immune Deficiency Syndrome
AF: Alexa Flour
CD: cluster of differentiation
DC: dendritic cel
dLNs: Draining Lymph Nodes
ELISA: Enzyme-Linked Immunosorbent Assay
FCS: fetal calf serum
FSC-A/H: forward scatter area/height
FcR: Fragment crystallisable receptor
GC: germinal center
gp: glycoprotein
HBSS: Hank’s balanced salt solution
HIV-1: human immunodeficiency virus 1
HLA-DR: human leukocyte antigen – DR isotype
HPLC: High performance Liquid Chromatography
i.d.: Intradermal
i.m.: Intramuscular
Ig: immunoglobulin
MHC-II: major histocompatibility complex I
MW: Molecular Weight
NHP: non-human primate
PBS: phosphate buffered saline
PMNs: polymorphonuclear neutrophils
RM: Rhesus macaques (Macaca mulatta)
ROS: reactive oxygen species
RPM: rounds per minute
RT: room temperature
RSV: respiratory syncytial virus
SSC-A: sideward scatter area

